# The external rumen of dung beetles: Complex interactions between larvae and their ontogenetic environments shape growth and life history

**DOI:** 10.1101/2025.03.21.644625

**Authors:** Nathan McConnell, Patrick Rohner

**Author notes:** Correspondence: Nathan McConnell, Department of Ecology, Behavior, and Evolution, University of California San Diego, 9500 Gilman Drive La Jolla, CA 92093.

## Abstract

Organisms are not just passive recipients of environmental pressures but are able to shape the environment they experience. However, the mechanisms and the evolutionary implications of such niche construction remains poorly understood. Larvae of the gazelle dung beetle (*Digitonthophagus gazella*) extensively modify their environment and benefit from microbial symbionts to digest their cellulose-rich diet. These modifications are so extensive that previous research suggests that dung beetle larvae establish an “external rumen”, where behavioral adaptations promote beneficial symbionts that enhance nutrient availability in the developmental environment. However, the mechanisms underlying these environmental modifications and their impact on species differences remains unclear. To investigate the external rumen hypothesis, we study the impact of larval environmental modifications on adult life-history traits in the dung beetle *Digitonthophagus gazella*. We did this by transplanting eggs into modified and unmodified environments, whilst excluding maternally derived microbes. Additionally, we include a heterospecific (*Onthophagus binodis)* manipulated environment to investigate evolution of species-specific effects. Counter to expectations, we find larval modifications by conspecifics did not confer a benefit to *D. gazella* in any aspect measured. However, surprisingly, focals from heterospecific treatments emerged significantly quicker. Additionally, we highlight the primary condition of the developmental environment as an essential factor in determining fitness benefits compared to any additive environmental effects. Our research adds to the growing literature on organism by environment interactions and demonstrates the relationship between dung beetle larvae and their developmental environment are complex and are not consistent with the presence of a simple external rumen.

## Introduction

Natural selection acts on the fit between an organism’s phenotype and its environment. Understanding the relationship between organisms and their environment is thus fundamental to evolutionary biology and there is a wealth of literature documenting how the environment drives (adaptive) evolution (Wallace 1877; Endler 1977). However, while organisms are often seen as passive recipients of environmental selective pressures, many actively shape the way they interact with their surroundings in diverse ways. For instance, organisms can change their phenotype in response to environmental conditions through developmental plasticity. This allows organisms to track changing fitness optima due to changes in the environment (DeWitt et al., 2004; West-Eberhard, 2005; Ghalambor et al., 2007). Furthermore, many organisms can shape the environment they experience through behavioral or physiological mechanisms that directly modify their environment, a process known as niche construction (Odling-Smee et al., 2003). By changing the environment an organism experiences, niche construction can in turn mediate plastic responses of the organisms in the modified environment. Understanding how an organism shapes its environment and responds to new environmental conditions is thus a central question in evolutionary biology.

Environmental modifications are particularly important when conditions are challenging. This is especially true for many detritivores, which inhabit and consume nutritionally poor, complex, and hard-to-digest food sources (Hungate, 1984). Many invertebrate detritivores lack the enzymes needed to break down complex carbohydrates and instead rely on symbiotic microbial communities for digestion (e.g., Brune, 2014). Some species even create entire external rumens, where microbial communities predigest the food source. The reliable acquisition and maintenance of these symbionts are therefore crucial (Hammer et al., 2019; Jones et al., 2024). Given that detritivores play key roles in ecosystem functioning (e.g., nutrient cycling: Hättenschwiler et al., 2005; disease control: Miller et al., 1961; and bioturbation: McCary et al., 2021) understanding the mechanisms that mediate organism-environment interactions is essential.

Dung beetles are an important group of detritivores that provide vital ecosystem services in agricultural grasslands (Losey et al., 2006; Beynon et al., 2015). In tunneling dung beetles, females (sometimes with the help of a male) bury a dung mass, meticulously shape it into a so-called “brood ball,” and typically deposit a single egg (Hanski et al., 1991). Once the larva hatches, it feeds on the brood ball and rapidly increases its weight about 25-fold before undergoing complete metamorphosis into an adult. Because the larvae spend their entire juvenile period inside their brood ball feeding on their recalcitrant diet, the properties of the brood ball has major implications for larval growth, as well as adult life history and fitness. For instance, it is well documented that maternal provisioning in terms of brood ball size, quality, and burying depth underground affects offspring development time, adult size, and survival (Shafiei et al., 2001; Carter et al., 2020; Kirkpatrick et al., 2022). In addition, dung beetle larvae benefit from microbial symbionts to digest their cellulose-rich diet, and vertically transmitted symbionts are known to be important mediators of larval growth. During oviposition, females deposit a microbial inoculate into the brood ball (a so-called ‘pedestal’), a fecal deposit that contains an array of microbes found within her gut to aid digestion (Estes et al., 2013; Jones et al., 2024). The presence of the pedestal in the brood ball aids growth and development rate of the larva (Schwab et al., 2016), suggesting that there are fitness effects. Interestingly, the phenotypic consequences caused by the experimental manipulation of maternal or larval behavior differ between species (Schwab et al., 2017), populations (Dury et al., 2020), and even among half-sib families within populations (Rohner et al., 2024b). This suggests that the reliance on (or capacity for) these environment-modifying behaviors can evolve.

Throughout larval development, larvae eat and excrete the same material repeatedly and work themselves through their brood ball multiple times. Schwab et al., (2017) demonstrated that this larval behavior enriches microbial communities with taxa able to digest complex carbohydrates. This could explain why larvae that are repeatedly placed into a new brood ball (and thus are exposed to much more fresh dung) still grow slower and emerge later. These findings suggest that niche construction behavior might benefit larval growth through changes in the microbial community and that beetle larvae effectively turn their brood ball into an ‘external rumen’ (see e.g., Swift et al., 1979) where microbial communities (pre)digest complex carbohydrates, making them available to the larva. Similar external rumens have been found between fungi and Eurasian wood wasps (Thompson et al., 2014), Bess beetles (Ulyshen, 2018), and leafcutter ants (Khadempour et al., 2016). It is thus likely that similar processes act in dung beetles, but the mechanisms that underly these brood ball modifications are still poorly understood.

Here, we explore components of the external rumen hypothesis in the dung beetle *Digitonthophagus gazella.* By explicitly excluding maternally derived microbes, we investigate how environmental modifications made by the larva feedback on larval development. To test whether there are beneficial effects of brood ball modifications, we expose newly hatched larvae to brood balls that were previously modified by a different larva. Based on the external rumen hypothesis, we expect that larvae introduced to a pre-modified brood ball will exhibit faster growth and development compared to those that must establish their own microbial community. Secondly, we explore whether the external rumen is species specific by introducing a *D. gazella* egg to a pre-constructed niche of a closely related dung beetle species (*Onthophagus binodis*), expecting larvae to perform better when their brood ball was previously inhabited by a member of their own species. As an additional measure, we use two separate controls to establish the effect of unmanipulated microbial communities. Taken together, our results indicate complex and unexpected relationships between larval niche construction and performance.

## Methods

### General laboratory populations

To investigate the external rumen hypothesis, we focus on the gazelle dung beetle, *Digitonthophagus gazella,* originally collected in Bastrop County, Texas. For comparison we also used *Onthophagus binodis,* originally from Waimea, Hawaii. These two species diverged about 40 million years ago (Parzer et al., 2018) but are ecologically similar. Both species are native to sub-Saharan Africa and overlap in their native and non-native ranges (Tyndale-Biscoe, 1990; Davis et al., 2020). Whilst *D. gazella* is found on a wider range animal dung, both species are consistently found in cattle (Davis et al., 2020). *O. binodis* is smaller in adult size compared to the focal *D. gazella*, however, the mean larval body weight over the first eight days shows some comparable ranges (Rohner et al., 2021).

### Experimental design

To test the effect of different brood ball conditions on larval growth of *D. gazella* (our focal species), we used four experimental treatments. Firstly, we placed *D. gazella* larvae into a brood ball that was previously inhabited by a conspecific for 8 days (*conspecific brood ball,* CON). The eight-day standard was determined by intersecting larval weight between both species (Rohner et al., 2021), whilst maximizing the time window for microbial growth. To test for species differences in the effect of brood ball modifications on larval performance, we also placed *D. gazella* larvae in to brood balls previously inhabited by *O. binodis* (*heterospecific brood ball,* HET). To account for changes in brood ball quality that may occur over time, we also exposed larvae to a brood ball that was left empty but was incubated in the same way as the heterospecific and conspecific brood balls (*8-day old unmodified brood ball (8-UB)*). To test how this incubation affected brood ball quality (desiccation, decay, microbial community changes), we also raised larvae in freshly defrosted brood balls (*fresh unmodified brood ball (1-UB)*). Below, we elaborate on the specific methodology.

To generate the individuals needed to prepare the conspecific and heterospecific brood ball treatments, we removed ten mature females of *D. gazella* and *O. binodis* from the general population and placed them in two circular 235mm x 210mm ovipositing chambers. The ovipositing chambers contained a compact mixture of sterilized soil and sand and topped with previously frozen cow dung (approx. 250g) and were kept in a controlled environment at 27°C and 50% humidity for six days. After six days all brood balls were carefully extracted by sifting the soil from the ovipositing chamber. Eggs were extracted from the brood-ball using sterile featherweight forceps and placed on a sterilized gauze mesh. The egg was then washed with a sterilization liquid (48.5ml dH20, 0.5ml bleach, 0.05ml Triton X; see Rohner et al., 2024b; Schwab et al., 2016) and distilled water to exclude the maternal microbiome and added to a 3.2g artificial brood-ball housed in a sterile 12-well plate (VWR 734-2324). Artificial brood-balls were made from sterilized cow manure that had been squeezed by hand through a cheese cloth, removing excess moisture to better replicate natural brood-balls (Shafiei et al., 2001). All plates were transferred into an incubator (Caron Insect Rearing Chamber) set at 27°C and 50% humidity. At the same time, a subset of artificial brood balls was left empty and also placed in the incubator (to generate incubated unmodified brood balls (8-UB)). All brood balls used to establish the conspecific, heterospecific, and incubated unmodified brood balls were incubated for eight days. Any larvae that inhabited the brood balls were removed on the eighth day. Brood balls within which eggs did not hatch or larvae did not survive until eight days were excluded from the experiment.

Two days into the initial setup of the experimental 12-well plates, four *D. gazella* ovipositing chambers were set up to generate focal individuals. Brood balls collected after six days. Focal eggs were extracted from the brood balls, washed with fresh sterilization liquid, and placed into conspecific (CON), heterospecific (HET), or incubated unmodified brood balls (8-UB) (after the larvae that previously inhabited these brood balls were removed). We also placed one fourth of all larvae into a fresh unmodified brood ball which consisted of a standard 12-well plate containing artificial brood balls (3.2g) defrosted 24 hr prior (1-UB). Newly deposited eggs hatch approximately after 24 hr period. This was done to control any desiccation or decay effects that would naturally happen when a brood ball is incubated (8-UB).

Due to the relatively long development and low fecundity of dung beetles and logistical constraints, the experiment was repeated in three consecutive temporal blocks.

### Data collection

Eggs were checked daily to assess the potential effect of brood ball manipulation on egg hatching success and to control the precise age of larvae later in the experiment. To quantify larval growth rate, we recorded total larval body weight on day five and day seven post hatching for all surviving larvae. Larval weight was measured using a Mettler Toledo scale (Model AB135-S). Insect growth curves are intrinsically nonlinear and often complex. We therefore focused on the initial “free” period of larval growth which likely approximates the maximal physiologically realized growth rate (Rohner et al., 2021). As an estimate of larval growth, we calculated instantaneous relative growth rates (RGR) following (Tammaru et al., 2007):

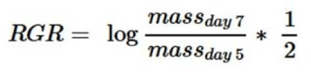

This estimate better approximates larval growth dynamics in a growing larva than “integral” estimates of growth rate that divide adult size by egg-to-adult development time (Tammaru et al., 2007).

Daily mortality checks were conducted until all individuals had emerged from the pupal stage, and adults were collected three days after emergence to allow cuticular hardening. The widest basal point of the pronotum was used as a proxy for body size, as established in dung beetle literature (Rohner et al., 2023). Pictures of the pronotum were obtained with a Pixelink camera (M20C-CYL) attached to a LEICA M205 microscope and measured in ImageJ (Version 1.54m) (Schneider et al., 2012).

### Statistical analysis

All statistical analysis was performed using R Core team V-4.4.1 (2024) (Code provided in supplementary material). Egg hatching success and larval survival were recorded as binary variables and were consequently analyzed using generalized mixed effects models from the lme4 package with a binomial family function (Bates et al., 2015). Larval survival was determined as the time an egg hatched to emergence as an adult. The three separate temporal replicate blocks were analyzed simultaneously, with experimental block designated as a random effect. Treatment was used as the only fixed effect.

Development time, relative growth rate (RGR) and pronotum width (adult size) were analyzed using mixed effect models from the lme4 package (Bates et al., 2015). Block was again added as a random effect. Treatment and sex were used as fixed effects. We always fitted full models, including the sex-by-treatment interaction but excluded the interaction term in cases where it was statistically not significant. For all models, Q-Q plots and histograms were used to check the residual distribution. A singular individual with an unusually large RGR had very strong impacts on the analysis and residual distribution and was therefore removed from the analysis. Where appropriate, differences between group treatments were analyzed using a Tukey post hoc test from the ‘emmeans’ package (Lenth, 2024) or from the ‘multcomp’ package (Hothorn et al., 2008). Effect size was calculated using partial eta-squared (η_p_^2^) values from the package ‘effectsize’ (Ben Shachar et al., 2020).

## Results

Exposing a developing larva to brood balls that were modified in different ways, we assess the degree to which larval brood ball modifications feedback on larval growth and performance. Neither egg hatch rates (Fig. S1) nor larval survival (Fig. S2) varied between the four treatments (hatching success: χ^2^ = 7.008, df = 3, p = 0.072, n = 217; larval survival: χ^2^ = 4.904, df = 3, p = 0.179, n = 161), suggesting that brood ball manipulations do not have immediate effects on mortality during early life stage transitions. However, egg-to-adult development time differed significantly by treatment (χ^2^ = 50.053, df = 3, p < 0.001, Fig.1A)). The effect size (η_p_^2^= 0.50, 95% CI: [0.31, 1.00]) suggests treatment explains 50% of the variation in development time. Specifically, larvae developing in a brood ball that was previously inhabited by a conspecific took longer to develop than larvae inhabiting a fresh unmodified brood ball (estimate = 4.93 ± 1.00 days, p <.001). Interestingly, larvae inhabiting a brood ball that was inhabited by a conspecific took marginally longer to reach the adult stage compared to larvae in brood balls that were modified by a heterospecific (estimate = 3.01 ± 1.16 days, p = 0.057). In contrast to our expectation, larvae inhabiting a fresh unmodified brood ball had significantly shorter development times compared to larvae placed into brood balls that were left empty for 8 days (estimate = 5.46 ± 1.03 days, p < 0.001) or inhabited by a conspecific (estimate = 3.54 ± 1.17 days, p = 0.020). We found no significant effects of sex (χ^2^ = 0.040, df = 1, p = 0.841) or evidence of an interaction between treatment and sex (χ^2^ = 3.566, df = 3, p = 0.312).

**Fig. 1.**
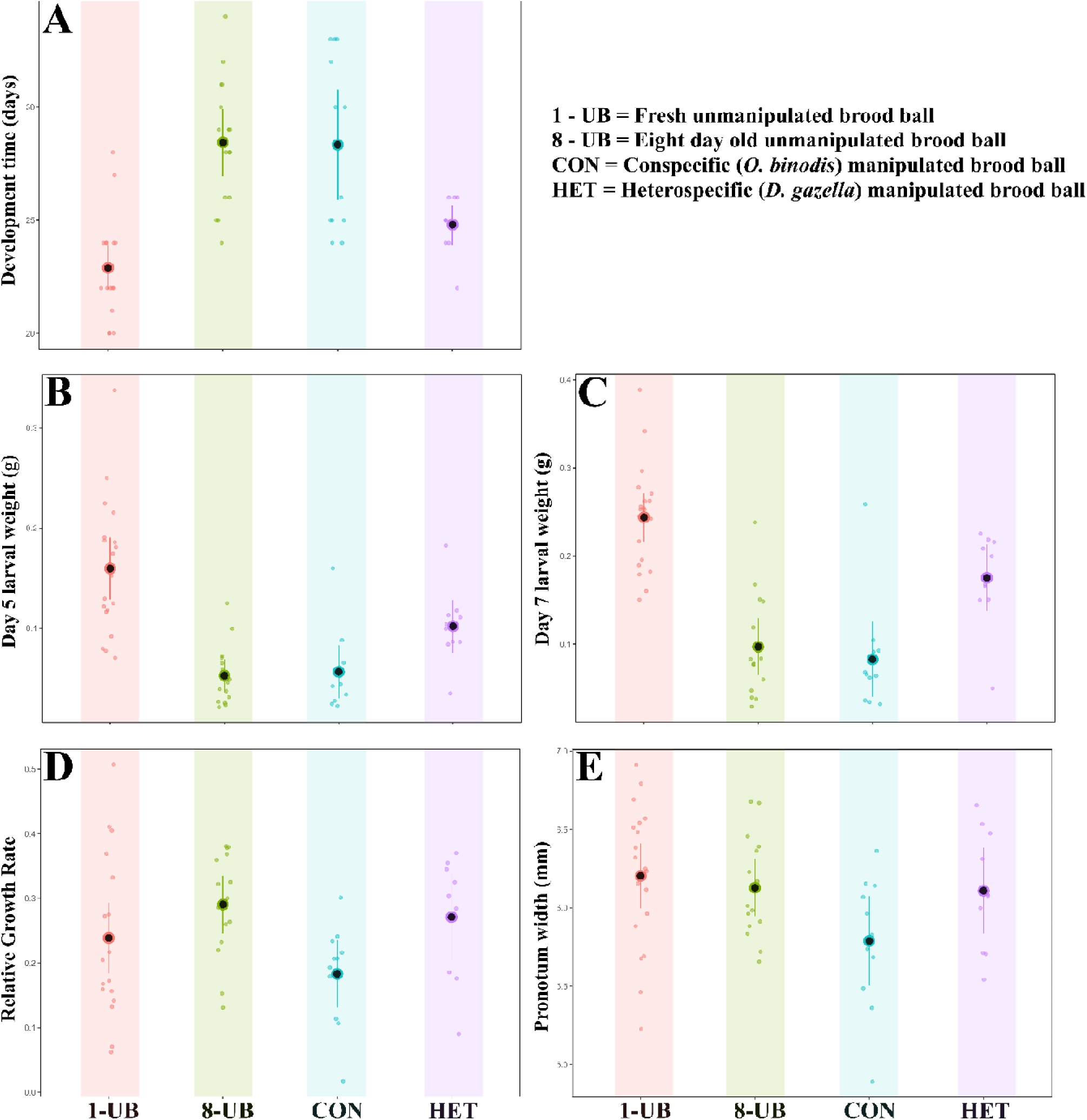
Developmental data for experimental treatments; Fresh Unmodified Brood Ball (1-UB), Eight-day old Unmodified Brood Ball (8-UB), Conspecific (CON) and Heterospecific (HET). Plots show means (Black dots) and corresponding 95% confidence limits. Colored points indicate treatment-specific individuals.

Larval weight at day five was significantly different by treatment (χ^2^ = 52.676, df = 3, p = < 0.001), but not by sex (χ^2^ = 0.217, df = 1, p = 0.641). The effect size (η_p_^2^= 0.50, 95% CI: [0.33, 1.00]) indicates treatment accounts for 50% of the variation we see. Pairwise comparisons show that specifically individuals in a fresh unmodified brood ball (1-UB) were heavier than larvae in an incubated, unmodified(8-UB) (estimate = 0.107 ± 0.019g, p = < 0.001), conspecific (CON) (estimate = 0.103 ± 0.018g, p = < .001), or heterospecific (HET) (estimate = 0.576 ± 0.019g, p = 0.019) brood ball. This demonstrates the importance of the larval environment in the early days of development.

Larval weight at day seven followed a similar trend to those at day five, with a significant effect of treatment (χ^2^ = 76.565, df = 3, p = <0.001), but not by sex (χ^2^ = 0.042, df = 1, p = 0.836). The effect size (η_p_^2^ = 0.68, 95% CI: [0.51, 1.00]) similarly shows the variation in weight explained by treatment is substantial (68%). Pairwise comparisons reveal that larvae in fresh unmodified brood balls (1-UB) continued to be significantly heavier than those reared in incubated unmodified (estimate = 0.147 ± 0.023, p = <0.001), conspecific (estimate = 0.160 ± 0.022, p = <0.001), and heterospecific (estimate = 0.069 ± 0.023, p = <0.018) brood balls. Interestingly, here the pairwise comparisons additionally show focal larva developing in brood balls modified by *O. binodis* (HET) were larger than focal larvae in brood balls modified by conspecifics (CON) (estimate = 0.091 ± 0.026, p = 0.005) and those in incubated, unmodified brood balls (8-UB) (estimate = 0.078 ± 0.026, p = 0.025). This further emphasizes the influence that the early environment has on dung beetle development time.

Relative larval growth rate showed a significant effect of treatment (χ^2^ = 7.839, df = 3, p= 0.049). However, effect sizes were low and reported large confidence intervals (η_p_^2^= 0.13, 95% CI: [0.00, 1.00]). Pronotum width (Fig. 1E) as a measure of body size was again significantly different by treatment (χ^2^ = 7.986, df = 3, p= 0.046), but not sex (χ^2^ = 2.135, df = 1, p= 0.144) or interaction between treatment and sex (χ^2^ = 2.49, df = 3, p= 0.477). Pairwise post hoc comparisons reveal individuals that developed in fresh unmodified brood balls (1-UB) were on average larger than individuals in conspecific brood balls (estimate = 0.407 ± 0.15, p = 0.043). As with relative growth rate, the confidence intervals of effect size are large (η_p_^2^= 0.15, 95% CI: [0.00, 1.00]). Collectively, these results show that larval brood ball modifications do not universally enhance larval growth. Instead, the effect of larval modifications emerge as context-dependent and only partially consistent with the external rumen hypothesis.

## Discussion

Dung beetles, like many other organisms that digest complex carbohydrates, rely on their microbiome to extract nutritional value from nutrient-poor substrates (Hungate, 1984; Parker et al., 2019). Previous research suggested the existence of an external rumen, where larvae actively manipulate and reshape the microbiome in their brood ball to benefit their growth. Here, we tested whether larval behavior contributes to an external rumen while excluding the confounding effect of maternally-inherited microbiomes and whether this is species-specific. In contrast to our expectations, we found mixed evidence for an effective construction of an external rumen. Irrespective of which variable investigated, *D. gazella* inhabiting a conspecific’s brood ball never improved performance compared to inhabiting an old brood ball. Instead, we recovered an unexpected pattern across species, suggesting more complex evolutionary and ecological interactions between larvae and their ontogenetic environment.

### Limited evidence of an external rumen in the absence of a maternal microbiome in D. gazella

Dung beetle larvae are known to alternate between eating the brood ball and redigesting their frass (Hanski et al., 1991) and drastically restructuring their brood ball throughout their juvenile development, leading to changes in the microbial capacity in degrading complex compounds (Schwab et al., 2017). These observations are consistent with the idea of an external rumen. However, if such behaviors were to establish an external rumen, we expected larvae to grow the fastest when placed in brood balls that had previously been inhabited by another larva. In contrast, the growth of larvae in pre-modified conspecific brood balls was indistinguishable from the growth of larvae in empty brood balls that were incubated for the same amount of time but without a larva. Surprisingly, the weight at days 5 and 7 as well as development time of larvae in newly defrosted brood balls was faster than both of these treatments, despite having no previous larval modifications. This also demonstrates that the early larval environment has a more significant impact on development time than any accrual of beneficial adaptations to the microbiome that may occur later. For example, the advantage of inhabiting a newly defrosted brood ball over an old unmodified one is to emerge as an adult almost five days earlier, and that is seemingly determined in those initial first few five days. Taken together, these findings suggest larval niche construction in *D. gazella* to be at best ineffectual, whilst in some instances detrimental, and ultimately, inconsistent with an external rumen.

### Potential evolution of larval brood ball manipulation

Organism-environment interactions are often taxon-specific (Holt et al., 2024; Schwab et al., 2017) and one would expect larvae to perform best in a brood ball that was modified by a conspecific. However, surprisingly, *D. gazella* larvae emerging from a conspecific’s brood balls did so at a significantly delayed rate compared to a brood ball previously inhabited by *O. binodis*. Larvae in the heterospecific brood balls were much heavier by day seven and, although not significant in pairwise comparisons, similar trends were found for growth rate and adult body size. Overall, this indicates that environmental manipulation can be beneficial, at least in some circumstances, and that these effects evolve across species.

The mechanisms underpinning these species-specific effects are, however, unclear. For instance, *O. binodis* may be more capable of developing or are better equipped in constructing a beneficial niche than *D. gazella*. Alternatively, the costs of inhabiting a conspecific’s brood ball could be explained by the initial *D. gazella* consuming a significantly larger proportion of the brood ball, thus, leaving the second (focal) larva with less available resources. It has been demonstrated that poorly provisioned brood balls directly impact the size of emerging dung beetles (Shafiei et al., 2001). However, the initial artificial brood ball was provisioned with 3.2g of dung, which is much more than required to reach maximum size (Rohner et al., 2021). Additionally, the eight-day standard was determined by intersecting larval weight between both species (Rohner et al., 2021), whilst maximizing the time window for microbial growth in the brood ball. Visual inspection of the non-focal *D. gazella* larvae did appear larger than *O. binodis* after eight days. However, the amount of dung available to the newly introduced focal egg did not seem significantly deficient, though it was more fibrous (note that all treatments were maintained under 27°C and 50% humidity). The observed differences may indicate that aspects of larval brood ball modifications evolve. On one hand, *O. binodis* might simply be better modifiers of an external rumen than *D. gazella*. Alternatively, the two species might differ in their target resources in early development, resulting in a depletion of specific elements established within the brood balls of a conspecific versus a heterospecific. Whilst this is speculative, further experimental designs should keep this aspect in mind in order to elaborate on the mechanisms driving these complex patterns.

It is important to consider that we here focus exclusively on the impact of larval behavior on brood ball quality while explicitly excluding parental effects, especially the maternally inherited microbiome. Previous studies indicate that the two factors are interacting in complex ways and this might explain the lack of the expected patterns in our study (Rohner et al., 2023; Rohner et al., 2024a). However, the lack of support for a role of larval behavior in the construction of an external rumen is still surprising and suggests that there might be other mechanisms at play. The external rumen hypothesis highlights microbial effects in nutrient provisioning. However, the striking difference between the effects of control brood ball age on growth outputs suggests that, instead of benefiting the growth of beneficial microbes, larval modifications might suppress the growth of harmful microbial taxa. Recent work in *O. taurus* report that microbial communities shift most distinctly between egg and larval life stages (Jones et al., 2024). This may highlight the necessity of a maternal pedestal to facilitate an external rumen or alternatively demonstrate that larvae are suppressing certain microbial taxa. Incubated unmodified brood balls accumulated visible microbial communities, sometimes leading to a large aggregation of fungal fruiting bodies. This was neither seen in the freshly defrosted control, nor in brood balls that were inhabited by a larva, suggesting that larval activity might disrupt the establishment of particular microbial communities. The suppression of harmful microbes is widespread across taxa (e.g. Amphibians: Harris et al., 2006; Fish: Masso-Silva et al., 2014; Reptiles: Van Hoek, 2014). Previous work on Grahams’s Frog (*Odorrana graham*) has recovered more than 370 antimicrobial peptides produced in its skin (Li et al., 2007) and in birds; the European Hoopoe (*Upupa epops*) uses metabolites produced by a symbiotic bacteria living in their uropygial gland that inhibits the growth of harmful microbes (Martín-Vivaldi et al., 2009). In insects the larvae of the fruit fly *Drosophila melanogaster* suppress the specific filamentous fungi growth through communal aggregations (Trienens et al., 2020). Whilst both the larvae of coconut rhinoceros beetle (*Oryctes rhinoceros*) and black soldier fly produce antimicrobial peptides effective at suppressing harmful microbes (Yang et al., 1998, Zhang et al., 2022).

Alternatively, the uninhibited microbial growth in incubated brood balls may have exhausted the nutritional value of the dung, thus slowing larval development time. However, if nutrition was restricted then we might expect body size to be significantly smaller (Rohner et al., 2021). The function of preventing certain microbes from being established thus appears more likely.

### Conclusions

Previous studies suggested that dung beetle larvae construct an external rumen that benefits their growth (Schwab et al., 2016; Parker et al., 2019; Patrick T. Rohner et al., 2023; Rohner et al., 2024b). Our results suggest that larval niche construction is more complex than previously thought. *D. gazella* larvae did not benefit from larval modifications of a conspecific, but did so when placed in a brood ball that was modified by a different species. This was unexpected and suggests that the benefits might only be found in some species but not others, or that we don’t fully understand all the factors involved in modifying brood balls. Furthermore, we only find limited support for increased nutrient availability through larval modification. Instead, our results suggest that brood ball modifications might hinder the establishment of specific microbes. More generally, our study highlights the complex and unexpected interplay between developing organisms, their effects on the environment, and how modified environment feeds back on organism growth, life history, and fitness (Rohner et al., 2024a; Sultan, 2015). Rather than viewing the relationship between organisms and their environment as a linear, unidirectional process in which environmental pressures simply shape selective outcomes, our findings suggest a more dynamic and reciprocal interaction. Organisms do not merely respond to environmental challenges; instead, they actively reshape their surroundings, which in turn influences their own development and potentially that of future generations. Future research will be required to shed light on the detailed microbial and behavioral mechanisms mediating these effects in dung beetles.

## Acknowledgements

We would like to thank collaborators who helped with collection, husbandry and stimulating discussions around this manuscript including Ben Mathews, Michelle Herrera, Sarah Britton, Ebony Argaez and Avi Khanna.

## Competing interests

The authors declare no competing or financial interests.

## Data availability

Raw data and code available at DOI: 10.5061/dryad.z8w9ghxqv

## Notes

https://doi.org/10.5061/dryad.z8w9ghxqv

### Competing Interest Statement

The authors have declared no competing interest.

